# AgAnt: A computational tool to assess Agonist/Antagonist mode of interaction

**DOI:** 10.1101/2021.11.11.468208

**Authors:** Bhavay Aggarwal, Arjun Ray

## Abstract

Activity modulation of proteins is an essential biochemical process in cell. The interplay of the protein, as receptor, and it’s corresponding ligand dictates the functional effect. An agonist molecule when bound to a receptor produces a response within the cell while an antagonist will block the binding site/produce the opposite effect of that of an agonist. Complexity grows with scenarios where some ligands might act as an agonist in certain conditions while as an antagonist in others [1, 3]. It is imperative to decipher the receptor-ligand functional effect for understanding native biochemical processes as well as for drug discovery. Experimental activity determination is a time extensive process and computational solution towards prediction of activity specific to the receptor-ligand interaction would be of wide interest.

## Introduction

Studies to classify agonists and antagonists such as [12] and [8] use molecular descriptors to classify androgen receptor and SlitOR25 ligands respectively. [6] also used molecular descriptors and fingerprints to classify androgen receptor ligands. [2] followed a similar approach to classify 5-HT1A ligands. [10] implemented extended connectivity fingerprints and extracted descriptors from ligands to classify TNBC and GPCR ligands. [9] used images of 3D chemical structures to predict Progesterone Receptor antagonist activity. These studies have achieved good results using random forest and SVM based models with some implementations using deep neural networks but all current methods take into account a single receptor when predicting its agonists or antagonists. We believe that the activity of a complex after ligand binding is determined by both the ligand and the receptor and thus both should be featurized to make predictions.

Machine Learning helps to uncover underlying patterns between various classes. When dealing with machine learning on proteins, the accuracy of the representations determines the final performance of the model. [5] used a feature map based on residue-based features. [7] use the raw protein sequence as well the fingerprints of the drug targets to predict drug-protein interactions. [4] represent a protein as a graph with residues acting as nodes, edges represent the spatial relationship between them and use Graph Convolution Networks for classification. Studies such as [11] have further explored various methods adopted to solve protein-related machine learning problems. We used 3 different approaches for this work, which are: - Sequence-Based Models Physiochemical Properties Based Models Graph-Based Models

## Results

We created our dataset by filtering through RCSB for entries which matched our criteria and further refined the results by performing additional filtering which is described in the supplementary section.

Models based on protein sequence performed the best on our experiments. Using Word2Vec, the protein sequences were represented as 100 length vectors. We experimented with different vector sizes but achieved better performance with length 100 as was also observed in other papers. Training these vectors without the ligands achieved 81% accuracy with 10-fold stratified cross validation. Using one-hot encoded SMILES representation as an additional input, we were able to bump up the accuracy by 4.6% with the best model performing at 86.5% accuracy (Fig. 1.1). Tree based models performed better than SVM’s which suggests non linearity in the ProtVec features. The feature importance plots of the AdaBoost and XGB classifiers also give further indication to their success. Out of 11500 features, only 47 were used by the AdaBoost model to make predictions. This suggests that by selecting these features, the model is able to get an advantage over SVM. Also, including one-hot encoded SMILES strings makes it difficult for SVM classifiers to handle such a large number of features. SMILES representations contribute to 32% of the features used by AdaBoost and the boost in performance is a confirmation of our pairwise approach to the problem.

**Figure 1:**
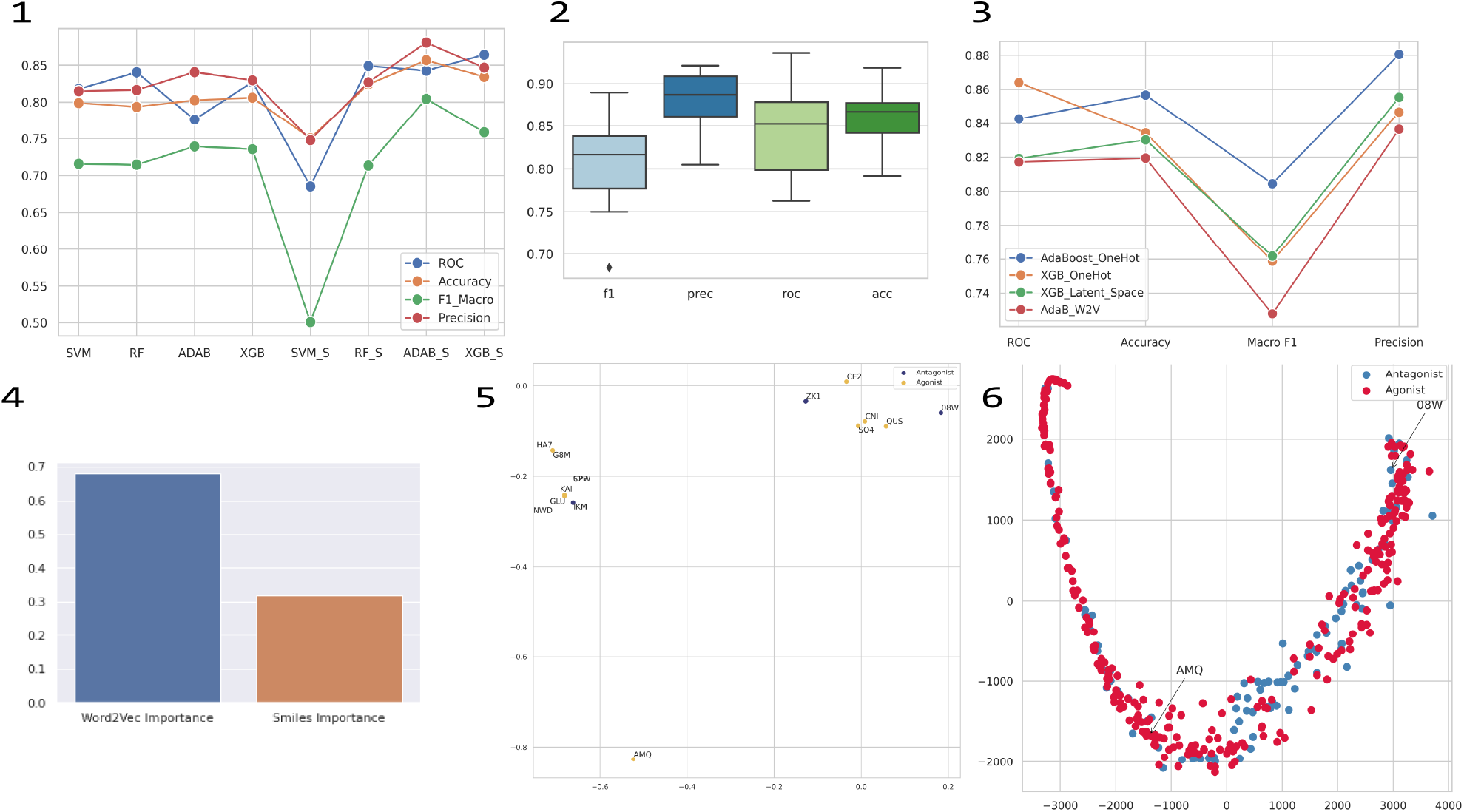
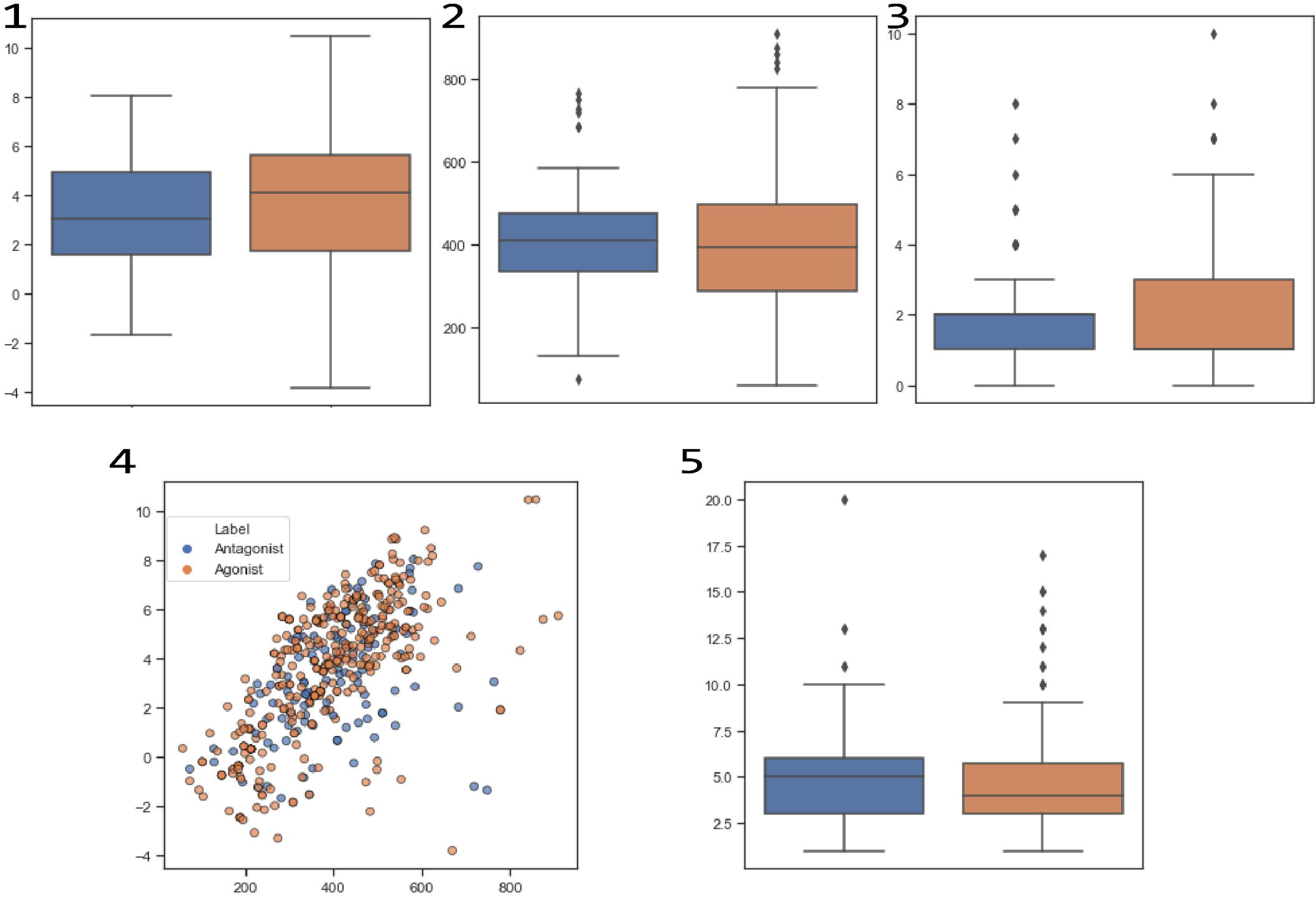
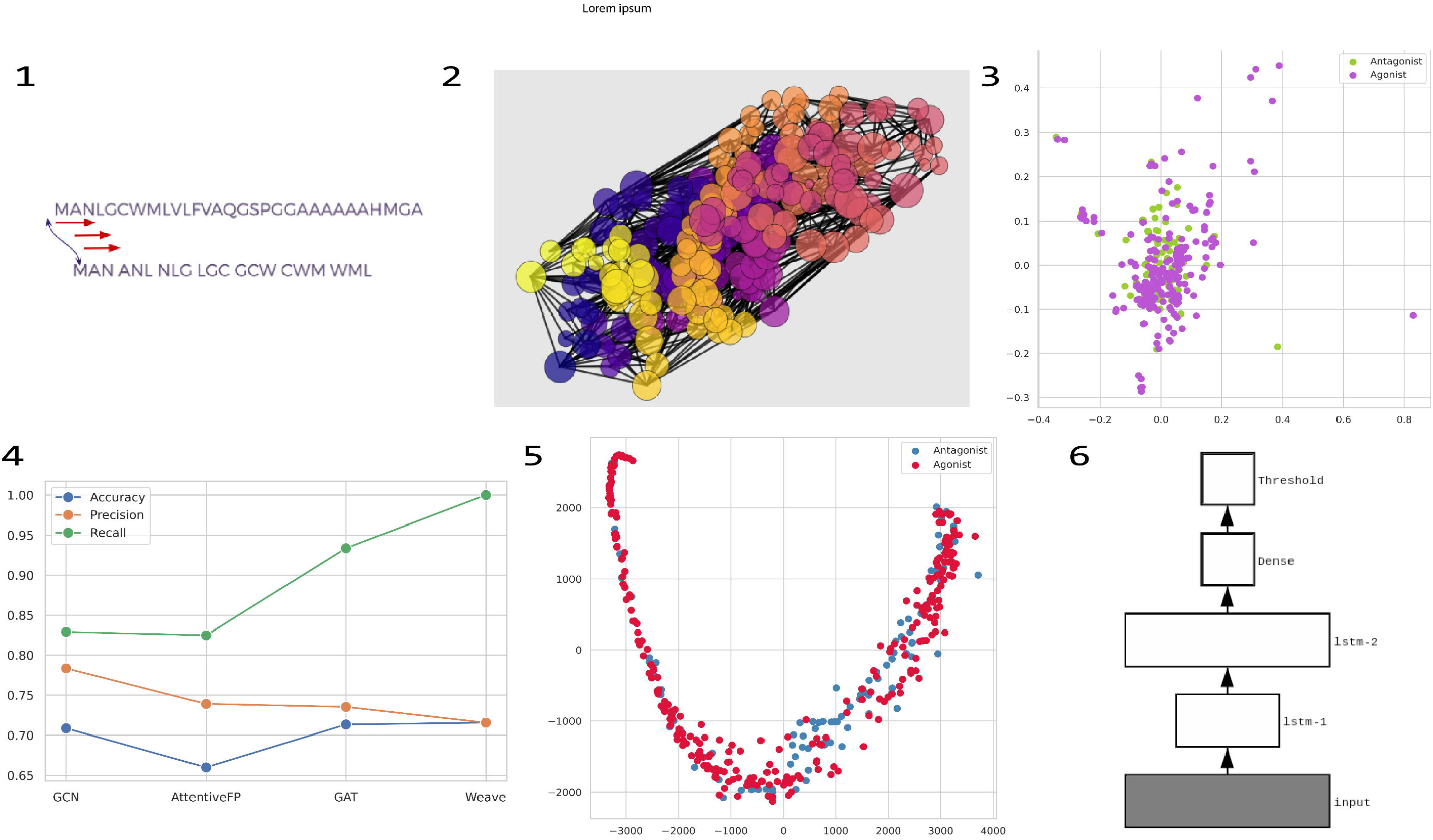
1) Results of the machine learning models trained on ProtVec (Model-Name) and ProtVec+SMILES features (Model-Name_S. 2) Cross validation scores of AdaBoost model trained on ProtVec+SMILES features. 3) Results of the best models trained of ProtVec + various SMILES representations. 4) Feature Importance Distribution for AdaBoost. 5) PCA of features showing only the ligands binding to Glutamate Receptor 6). PCA of ligand fingerprints highlighting ligands AMQ and 08W.

XGB Classifier performed best with Latent space embeddings while AdaBoost outperforms XGB slightly when using Word2Vec SMILES embeddings achieving an accuracy of 0.819 in both the cases. One-hot encoded representation of SMILES outperforms more efficient representations tested possibly owing to the inferences from our previous finding of only a few features being important. It seems that in the process of reducing dimensions for a more efficient representation, information important to our classification task is being lost. Also, we find that AdaBoost with one hot SMILES uses 46 (30 protein, 16 smiles) features whereas with Word2Vec embedded SMILES it only uses 42 (23 protein, 19 smiles) which is an unexpected decrease in the number of protein features being used by the model. Some important information of SMILES gets lost in these representations which is why one hot encoding performs better and although these representations are much more efficient, one hot encoding is more suitable for our classification task.

We created LSTM’s and Graph Neural Networks to utilize the physiochemical and structural properties of proteins. These models were not able to perform as well as our sequence based models and have been detailed in the supplementary section.

Further analysing our pairwise approach, we looked at the change in activity of a protein as the binding ligand changes. One such case was Glutamate Receptor 2. GLUR2 exhibits agonistic activity when ligand AMQ binds with it while exhibiting antagonistic activity when ligand 08W binds to it. When we consider all the ligands, their fingerprints dont show any seperation between agonists and antagonists. Same is the case with receptor features and we believe that it is the combination of the properties which allows our model to differentiate between agonists and antagonists. This is further indicated by Fig 1.5, fingerprints of AMQ and 08W when considered individually indicates distinction in physical and chemical properties. The properties of the two ligands in table X also indicates a difference in the two ligands. Similarly, ligand CNI exhibits agonistic activity when bound with GLUR2 but exhibits antagonistic activity when bound to GLUR3. This difference must arise from the difference of structure and properties of the two proteins in this case and all such cases. These distinctions only help us further our pairwise hypothesis and do not give much information on mechanisms which determine the activity upon binding of the ligand.

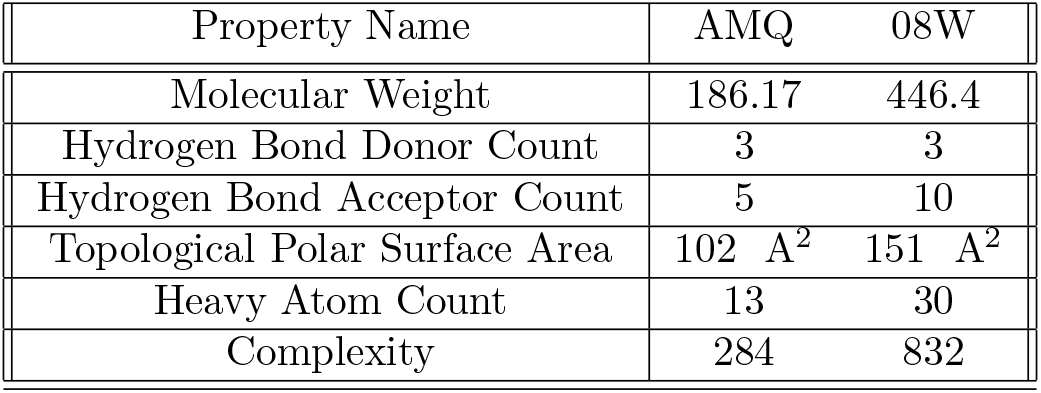

| Property Name | AMQ | 08W |
| --- | --- | --- |
| Molecular Weight | 186.17 | 446.4 |
| Hydrogen Bond Donor Count | 3 | 3 |
| Hydrogen Bond Acceptor Count | 5 | 10 |
| Topological Polar Surface Area | 102 A<sup>2</sup> | 151 A<sup>2</sup> |
| Heavy Atom Count | 13 | 30 |
| Complexity | 284 | 832 |

Our model is hosted on the domain agant.raylab.iiitd.edu.in and can make predictions for any given PDB ID-Ligand ID pair or Protein Sequence-Ligand SMILES pair. Additional usage instructions are detailed on the web page. The scipts for this project are available on the [Github](https://github.com/RayLabIIITD/AgAnt) repository.

## Discussion

Advances in Natural Language Processing models and the accomplishments of models like GPT-3 in producing text incredibly similar to humans shows that machine learning models have learnt to capture the underlying principles of language. Creation of similar models specifically for proteins will enhance all kinds of machine learning tasks related to proteins.To achieve this, we believe that there is a need to better represent protein features and an efficient integration of structural features with protein sequence. We explored many possible approaches which lays some groundwork for our future experiments. ProtVec despite being a relatively simple approach produces strong results though has problems with generalization and might not be suitable for larger datasets. ProtTrans assigned tokens to each residue and extensions to this approach might lead to better learning techniques. Physiochemical features of residues performed well though their performance on larger datasets and more advanced models are some things that still to be experimented with.

Graphs still are the most visually apt representations of proteins but require better features in order to solve such classification problems. Entire graphs although give an accurate representation of protein structures, are harder to extract features from given their size. Protein graphs show properties similar to images,-

Locality - nearby residues are alike and as distance increases, dissimilarity increases. Stationarity - features can appear anywhere in the graph. Compositionality - features follow a hierarchy.

Which is why we believe that GCN’s can achieve much better results as CNN’s have on image tasks. We believe that these representations should be used only for much larger datasets so effectively utilize the convolutional networks.

## Conclusion

In summary, we have demonstrated our models ability to differentiate between agonist and antagonist protein-ligand pairs with high accuracy. We have examined various representations of proteins and different machine learning models that can be used for classification problems. AdaBoost model trained on ProtVec and SMILES representations was the most precise and accurate model. Our results demonstrate that featurizing both the ligand and protein is not only theoretically accurate, it experimentally performs better than only considering a single entity. In the future, we want to experiment with better representations of proteins on specialized language models. The success of models based only the sequence of the protein show potential for the application of specialized transformer models for proteins.

