## Supplementary Document for "AgAnt: A computational tool to assess Agonist/Antagonist mode of interaction"

### Materials and Methods

**Dataset:** To collect the relevant protein files, RCSB[?] database was used. All ID's matching with the search terms " Protease bound with agonist " and "Protease bound with antagonist " were extracted. Files containing RNA/DNA were removed followed by automated filtering where files which did not contain any words Agonist/s or Antagonist/s in their PDB title, header or its primary PubMed citation were removed. Files were then manually filtered using the above-mentioned criteria and also, on the ligands and molecules present in the file. Negative ID's and duplicates were removed and the final data composition was as follows –

551 Proteins 156 Antagonists 395 Agonists

Peptides ligands were also removed when considering the ligands as features in the models which left 489 proteins in which 131 were antagonists and 358 agonists. Additionally, there are 214 unique proteins and 423 unique ligands. We performed exploratory analysis on these ligands to investigate whether there is any distinction between ligands acting as agonist or antagonist. We extracted Lipinski descriptors using rdkit and visualized their distribution. Also, pubchem fingerprints of the ligands were extracted using PyFingerprint[?] and to visualize the two classes, PCA was used.

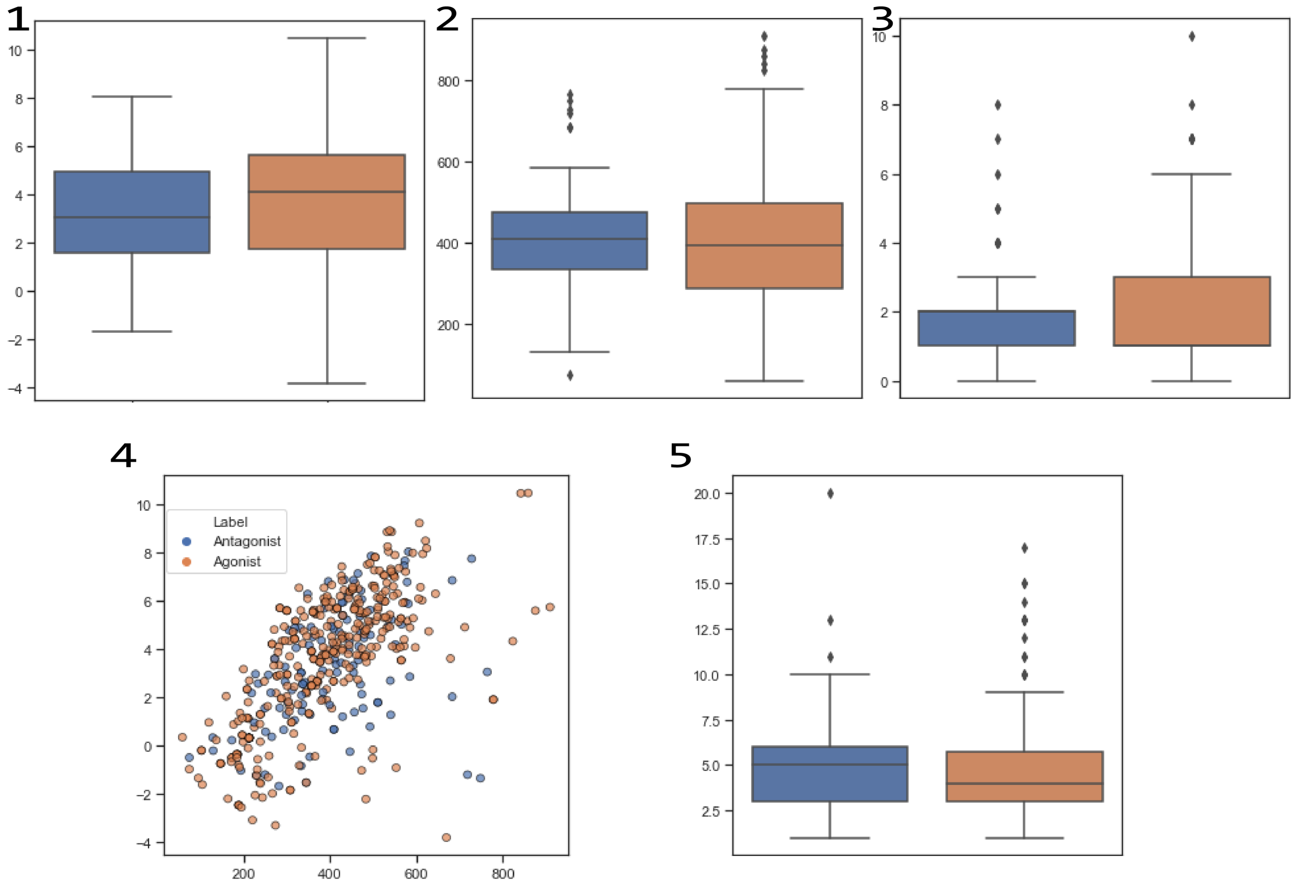


Figure 1: 1) Boxplot for Octanol-Water Partition Coefficient (LogP) of the ligands. 2) Boxplot for Molecular Weight (MW) of the ligands. 3) Boxplot for Number of H-Donors in the ligands. 4) Scatterplot of Molecular Weight vs Octanol-Water Partition Coefficient (LogP) of the ligands. 5) Boxplot for Number of H-Acceptors in the ligands.

The plots in Figure 1 indicate that there is no significant difference between ligands acting as agonist and antagonist. Each feature indicates no significant difference which is also the case with PCA on ProtVec representations which can be seen in Fig 2.5.

**Sequence-Based Models:** In this approach, we only consider protein sequence while making predictions. We chose to baseline this approach with ProtVec[?] which is a common yet powerful way to generate embeddings. ProtVec applies Word2Vec which is a method to generate word embeddings. The word context is learnt and the embeddings are generated in such a way that positions similar words closer in the vector space. For proteins, overlapping windows of residues of size 3 are considered as words. The embeddings created are summed up to create one vector to represent the protein. These embeddings were then trained on machine learning models with cross-validation. To handle the class bias, we used Stratified k-folds which mains the class ratio in folds as it is in the original dataset. Models were trained using sklearn and xgboost. The default parameters were used for reproducibility and the cross-validation was done using sklearn with 10 as the number of folds.

Up till now, the ligand has not been taken into account while making predictions. To incorporate ligands, we use their SMILES representations and one hot encode them. These embeddings along with the previously generated ProtVec representation were combined to generate representations for the protein.

Other methods to encode SMILES were also used namely SMILESVec[?] and ChemVAE[?] SMILESVec encodes SMILES using Word2Vec whereas ChemVAE uses variational autoencoders to generate latent space embeddings. Both these methods produce embeddings which have lower dimensions compared to one-hot encoding.

We also experimented with state of the art Natural Language Processing models namely BERT[?] , RoBERTa[?] and ELECTRA[?] . We first started with NBSVM - Naive Bayes Support Vector Machine which is a baseline language model. The protein sequences were processed into sequences of overlapping windows which is how they were used in ProtVec. This method trains a support vector machine using naive Bayes log-counts asfeatures. We were able to achieve 76% validation accuracy but the test accuracy was barely 55%. We decided to move on to more advanced language models and trained BERT, RoBERTa and ELECTRA from scratch. BERT - Bidirectional Encoder Representations from Transformers, is a bidirectional training approach which applies attention mechanism to language modelling. This helps better capture the contextual relation between words, and its bidirectional architecture allows it to consider the surroundings on both sides. RoBERTa builds upon BERT by tuning its training procedure. ELECTRA applies pretraining to BERT called replaced token detection. This helps the training process especially for smaller models. Their vocabulary was trained on all possible 3-length amino acid sequences. The protein sequences were split into overlapping windows as it was done for ProtVec. The models when trained on our dataset would only predict the majority class agonist. This model behaviour continued even when the model was trained on 10k randomsequences and then fine tuned on our dataset. This might be because these models expect sequence length to be a maximum of 512 but our sequences range upto 10k words. Also, our method of generating sentences -length 3 overlapping windows, might not be a good suit for such methods. To further investigate this, we used ProtTrans [7] to generate embeddings for our protein sequences. ProtTrans has language models designed forproteins and are trained on millions of protein sequences. Using ProtBert-BFD, we generated embeddings forour protein sequences and trained them on AdaBoost which performed best using ProtVec. Instead of dividingthe sequences into overlapping windows, ProtTrans assigns each amino acid an operate token and supportslarger sequence lengths - 40k for the model we chose. ProtTrans performs comparatively better than similartransformers but classic methods like ProtVec still outperform for our classification task. This result indicatesthat transformers specialised for proteins have potential to produce good results but require a larger datasetand downstream fine tuning for specific classification tasks.

**Physiochemical Properties Based Models:** The physicochemical characteristics, such as molecular size, net charge and amino acid composition are some of the factors that contribute to the functional properties of proteins. Compared to solely using the sequence, these features add additional information about the composition and various properties of the sequence but are extracted solely from the sequence of the protein and again do not use the structural information present in a PDB file.

iFeature[?] was used to extract features from the protein sequences. There were 20 features extracted and are described in the Supplementary,

Since these features are dependent on the protein length, they have different dimensions for different proteins. To deal with these variable-length features, we used RaggedTensors from Keras to train LSTM models. Apart from the features mentioned above, amino acid index features were extracted from AAindex. These features are 20 numeric values representing various physiochemical and biological properties of residues.

Long Short Term Memory networks are recurrent neural networks with a feedback mechanism which helps maintain the gradient flow. This is of interest to us because residue interactions range from short to long distance and to capture these effectively, LSTM's overcome the short-term memory issue in RNN's by identifying important parts of the sequence and passing them downstream. This enables LSTM's to learn over entire sequences and identify regions of interest which improves learning on sequential data. We trained multi layer LSTM and Bidirectional LSTM model. Cross-validation with stratified k folds from sklearn was again used to make generalized predictions.

Using Optuna, we optimized our hyperparameters of our LSTM model for different architectures. The best architecture was a double layer LSTM followed by a Dense layer as visualized in (Fig. 2.6). The model was then trained with stratified 10 fold cross validation and the results are shown in. Other architectures had similar scores and performed well for the size of the data present. Addition of features namely AAIndex and ProtVec sequences led to a decrease in performance. Our results show that, addition AAIndex features did not have any positive impact on the models performance which we hypothesize is because they are just positional replacements of amino acids with physicochemical feature values. The performance decrease on the addition of ProtVec sequences can be attributed to the loss of sequence properties in the process of creating Word2Vec embeddings apart from the issues such as the size of the database which is not suitable for deep learning models. This approach had slightly better performance than vanilla but would require a much larger dataset to fully utilize the capabilities of LSTM's. The inclusion of ligand features might also boost the models performance.

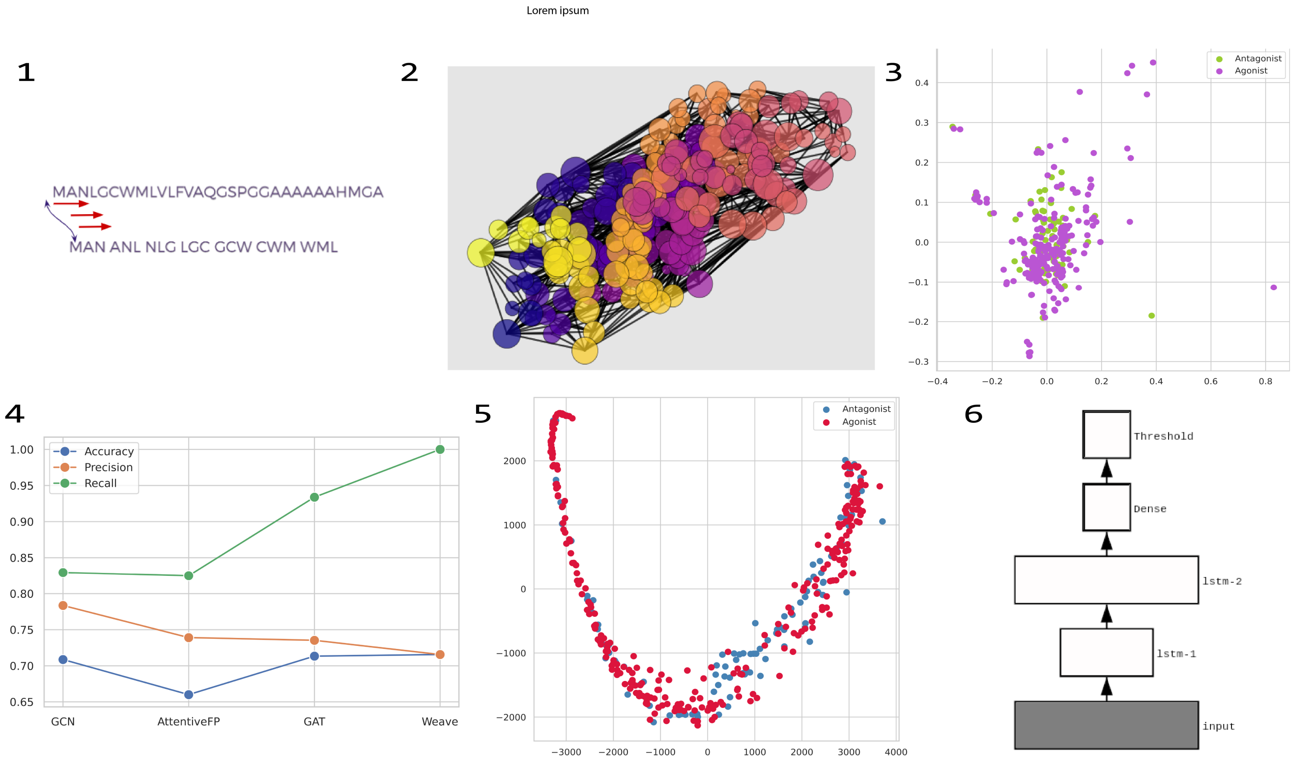


Figure 2: 1) Representation of ProtVec preprocessing of the protein sequence. We experimented with window sizes 5 and 7 but size 3 outperformed both. Embedding size was chosen to be 100. Other sizes performed worse. 2) Graph generated by Graphein of protein with ID 6HLL. 3) PCA on ProtVec features of the ligands. 4) Result of experiments using Graph neural Networks. 5) PCA on PubChem Fingerprints of the ligands. 6) Representation of our LSTM model.

**Graph-Based Models:** Graphs are structures which represent the relation between objects. Representing atoms as nodes and their bonds as edges, it is a far more accurate representation of chemical molecules when compared to matrices generated to represent features. To incorporate the structural properties of proteins, it is imperative to featurize their spatial arrangement. This would help to include the residue interactions, cavities and other properties. To generate these graphs, we used Graphein[?] as visualized in Fig 2.2. The graphs capture the internal chemistry of proteins and featurizes residue features as node attributes while bonds get representation of edges. The coordinates of residues are preserved which helps the model learn structural properties. Node features include amino acid embeddings, secondary structure assignments, solvent accessibility and the coordinates. Edges contain features describing distances. We generated graphs for all the PDB files and used DGL[?] to train Graph Neural Networks. We used 4 types of GNN's -

- Graph Convolutional Network (GCN) - graph counterpart of convolution neural networks, convolutions help learn features from neighbouring nodes.
- Graph Attention Network (GAT) - extends GCN architecture to apply attention among node neighbourhoods.
- Graph Attention Network (AttentiveFP) - graph attention mechanism based neural network.
- Molecular graph convolution (Weave) - graph convolution with the addition of weave modules which help produce better representations of the input.

| Model Name | Accuracy | Precision | Macro_F1 |
|---|---|---|---|
| NBSVM | 0.50 | 0.25 | 0.33 |
| Transformers (BERT, RoBERTa and ELECTRA) | 0.48 | 0.48 | 0.33 |
| ProtBert-BFD on AdaBoost | 0.70 | 0.77 | 0.60 |
| LSTM | 0.80 | 0.84 | 0.74 |
| GCN | 0.73 | 0.77 | 0.61 |

Graphs created by Graphein were again trained with 10 stratified fold cross validation. All the four modelshave similar performance with GCN performing the best in terms of predicting the minority class i.e antagonists.Weave only predicted the majority class whereas GAT and AttentiveFP were able to predict antagonists butwith poor precision. To overcome the training issues caused by our relatively small dataset for such a task, themodels were trained on 10k randomly chosen proteins. The classification layers were then removed from the trained model and the model was fine-tuned on our dataset. This had minimal impact on the performance of the models with GCN again performing the best but comparatively worse than other techniques and can be seen in Fig. 2.4 and Table .

**Metrics for evaluation of our models:**

Accuracy -

$$\frac{TP + TN}{TP + FP + FN + TN}$$

Precision -

$$\frac{TP}{TP + FP}$$

F1_Macro =

$$\frac{1}{N}\sum_{i}^{N} 2\frac{Precision_i * Recall_i}{Precision_i + Recall_i}$$

Where N is the number of classes, and $metric_i$ corresponds to the metric value for the ith class. Apart from accuracy, we want to maximize our F1 score to ensure that the model is predicting the minority class i.e. antagonist with good accuracy.

Apart from the above metrics, we also use ROC curves which are a visual representation of the performance of the model at different thresholds. It is quantified by the area under the curve.

Features extracted from IFeature -

Features extracted from AAIndex -

- Hydrophobicity index
- Residue volume
- Transfer free energy to surface
- Steric parameter
- Polarizability paramete
- The Chou-Fasman parameter of the coil conformation
- Localized electrical effect
- Average accessible surface area
- Positive charge
- Negative charge
- Buriability

| Feature Name | Description |
|---|---|
| AAC | Amino Acid Composition is the frequency of each amino acid in the protein sequence |
| CKSAAP | Composition of k-spaced Amino Acid Pairs is the frequency of amino acid pairs separated by any k residues |
| DPC | Di-Peptide Composition |
| DDE | Dipeptide Deviation from Expected Mean |
| TPC | Tri-Peptide Composition |
| GAAC | Grouped Amino Acid Composition further classifies amino acids into 5 classes |
| CKSAAGP | Composition of k-Spaced Amino Acid Group Pairs is the frequency of amino acid group pairs separated by any k residues |
| GTPC | Grouped Tri-Peptide Composition |
| NMBroto | Normalized Moreau-Broto Autocorrelation descriptors |
| Moran | Moran autocorrelation descriptors |
| Geary | Geary autocorrelation descriptors |
| CTDC | Composition descriptor is the composition of polar, neutral and hydrophobic residues of the protein |
| CTDT | Transition descriptor is the frequency with which a residue either polar, neutral or hydrophobic is followed by a residue of a different group |
| CTDD | Distribution descriptor describes the spread of the amino acid groups(polar, neutral or hydrophobic) across the sequence |
| CTriad | considers vicinal triad of amino acids as a single unit to determine properties |
| KSCTriad | extends CTriad to also consider units separated by k residues |
| SOCNumber | Sequence-Order-Coupling Number |
| QSOrder | Quasi-sequence-order |
| PAAC | Pseudo-Amino Acid Composition characterizes protein sequence using a matrix of amino-acid frequencies. Compared to AAC, additional information is also included such as the correlation between residues of a certain distance based on properties such as hydrophobicity value, hydrophilicity value and side-chain mass |
| APAAC | Amphiphilic Pseudo-Amino Acid Composition |
